# Uncovering the history of recombination and population structure in western Canadian stripe rust populations through mating-type alleles

**DOI:** 10.1101/2023.03.30.534825

**Authors:** Samuel Holden, Guus Bakkeren, John Hubensky, Ramandeep Bamrah, Mehrdad Abbasi, Dinah Qutob, Mei-Lan de Graaf, Sang Hu Kim, Hadley R. Kutcher, Brent D. McCallum, Harpinder S. Randhawa, Muhammad Iqbal, Keith Uloth, Rishi Burlakoti, Gurcharn S. Brar

## Abstract

The population structure of crop pathogens such as *Puccinia striiformis* f. sp. *tritici* (*Pst*); the cause of wheat stripe rust, is of interest to researchers looking to understand these pathogens on a molecular level, as well as those with an applied focus such as disease epidemiology. Cereal rusts can reproduce sexually or asexually, and the introduction of novel genetic lineages has the potential to cause serious epidemics such as the one caused by ‘Warrior’ lineage in Europe. In a global context, *Pst* lineages in Canada were not well-characterized and origin of foreign incursions was not known. We used a whole-genome/transcriptome sequencing approach for the Canadian *Pst* population to identify lineages in a global context, origin, and evidence for foreign incursion. More importantly, for the first time ever, we identified nine alleles of the homeodomain mating-type locus in the worldwide *Pst* population and show that previously identified lineages generally exhibit a single pair of these alleles. In addition, we find only two pheromone receptor alleles. We show that the recent population shift from the ‘*PstS1’* lineage to the ‘*PstS1-related’* lineage is also associated with the introduction of a novel mating-type allele (*b-3*) to the Canadian population. We also show evidence for high levels of mating-type diversity in samples associated with the Himalayan center of diversity for *Pst*, including a single Canadian race previously identified as ‘*PstPr’* (probable recombinant) which we identify as a foreign incursion from China circa. 2010. These data provide comprehensive details on the population biology of Canadian *Pst* diversity and mating-type alleles in the global *Pst* population which can be utilized in testing several research questions and hypotheses around sexuality and parasexuality in rust fungi.

## Introduction

*Puccinia striiformis* f. sp. *tritici*, the cause of stripe or yellow rust disease, is one of the five most important wheat pathogens in Canada and several epidemics of the disease impacted wheat production over last two decades [1]. Efforts to understand virulence phenotypes are ongoing [2–5] while genetic population structure studies [6,7] were limited in the Canadian landscape. Studying the genetic population structure of the pathogen in a global context is important because rust pathogen propagules can easily spread from one country to another with wind currents and even inter-continental spread in wheat rusts is reported [8]. Presence of foreign incursions of *P. striiformis* f. sp. *tritici* races in Canada has been speculated [7] but no study presented evidence of such incursions or information on origin of such incursions. While our research group studied the genetic population structure of Canadian *P. striiformis* f. sp. *tritici* populations, we utilized our generated and publicly available genomic resources to characterize mating-type alleles in the global pathogen populations. Mating-type alleles in wheat rust pathogens remained largely uncharacterized and no study showed utilization of mating-type alleles in answering biological questions relating to sexuality or population biology.

Mating in basidiomycete fungi such as rusts, smuts, and agaricomycotina depends upon a variety of factors including the development of sexual macrostructures at a certain life cycle stage, environmental cues, chemical signalling between individuals, and genetic compatibility. From a genetic perspective, non-self-recognition to facilitate mating in rusts is controlled by two unlinked loci: P/R (sometimes called *a* and equivalent to the *B* locus in agaricomycotina) and HD (sometimes called *b* and equivalent to the *A* locus in agaricomycotina). The P/R locus encodes pheromone precursors (*mfa*) and receptors (*Pra*) which must be compatible in order for prospective mates to signal to one another and initiate syngamy. The HD locus encodes two homeodomain genes (*bW-HD1* and *bE-HD2*) which need to be of different allelic specificity in each mate in order for their protein products to dimerise into heterodimeric bW/bE homeodomain transcription factors which regulate cellular development during mating, and for the rest of the fungus’ life-cycle including maintenance of the dikaryotic state and controlling pathogenicity in various smuts and rusts [9–11,16,17]. In some basidiomycota, the P/R and HD loci have become linked, leading to a bipolar rather than tetrapolar mating-type [12]. In other cases, the alleles no longer discriminate against self-fertilization or are not required for mating, leading to a bipolar or even unipolar mating-type [10]. So far, all characterized rust fungi are tetrapolar [18], although very little is known about the majority of rusts’ mating-type loci.

The biochemical and genetic mechanisms of pheromone signalling at the P/R locus are complex, and have been more extensively characterized in the Agaricomycotina and Ustilagomycotina [11,13–15]. A number of models for the locus in Puccinomycotina exist, most recently reviewed in [16], but overall: two discrete cells must carry complementary pheromones and receptors in order to initiate syngamy. In three wheat rust pathogens i.e., *P. triticina, P. graminis* f. sp. *tritici*, and *P. striiformis* f. sp *tritici*, three *Pra* receptor genes belonging to the STE3 family have been identified with *STE3.2-2* likely being a non-mating-type receptor, and *STE3.2-1* and *STE3.2-3* the likely mating-type pheromone receptors and hence nucleus / haplophase-specific [17]. Additional genes speculated to encode pheromone precursors (*mfa*) have been identified but none have been well-characterized, in part due to their short length and lack of conservation which makes them difficult to characterize. The three STE3 genes in different species consistently segregate when organized using phylogenetic methods, clustering with their orthologues and not their paralogues from the same species. Where sequence information from multiple isolates is available; additional STE3.2 alleles in rusts have not been identified, leading to the hypothesis that the three genes collectively comprise two complementary alleles, one of which is inherited with each haplotype. In this model, any dikaryotic rust cell will encode both *P/R* alleles, and haploid germ cells will have a 50% chance of being compatible with another germ cell.

The HD locus encodes a pair of homeodomain genes: *bW-HD1* and *bE-HD2* which are necessary for proper development of the fused dikaryon into a complete individual [11]. In *Pucciniales spp.*, the genes are ∼1200 and 1800 bp in length, with one and two introns respectively, encoding ∼400 and ∼600 amino acid length proteins [17]. Each protein exhibits three domains: at the N terminus a Variable domain, a central structured Homeodomain, and a Constant domain at the C terminus. bW-HD1 and bE-HD2 are entirely dissimilar on an amino-acid level, except for some (∼50% AA similarity) in the homeodomain region. Evidence from heterologous systems indicates that the two gene products physically associate in the cell and act via DNA-binding activities of the homeodomain [15,19,20]. The mechanism for non-self-recognition is that variable domains of the same mating-type prevent dimerization. Experiments with chimeric variable domains indicate that a relatively small alteration to the variable domain is enough to permit interaction [21–23]. Without the dimerized proteins, mating will not proceed as nuclei will not properly segregate into the daughter cells. Additionally, the normal cellular growth process cannot proceed without this same nuclear regulation. In *P. triticina*, at least nine mating-types have been identified (Guus Bakkeren, personal communication). Analysis of the published genome assemblies of *P. striiformis* f. sp. *tritici* identifies two alleles of each *Pst-bW-HD1* and *Pst-bE-HD2* in each genome on chromosome 4, although these are not always present in the published assembly, and five distinct alleles of each *Pst-bW-HD1* and *Pst-bE-HD2* in total, with the P/R locus being on chromosome 9 [18]. It has been hypothesized that maintenance of a plurality of mating-types is evolutionarily unfavourable without selection pressure to maintain outcrossing in sexual reproduction [10,24,25]. Loss of mating-type diversity through translocation to collapse HD and P/R into a single locus, as well as recombination events leading to self-fertility have both been observed in a number of related basidiomycete fungi [10,11], however, thus far the *Pucciniales* all exhibit maintenance of this system.

Given that the global *P. striiformis* f. sp. *tritici* populations in any particular place and time are often composed of a small number of asexually reproducing clades, often with one highly dominant lineage which is best adapted to local growing conditions [26], and also given that *P. striiformis* f. sp. *tritici* mating-types are expected to be reasonably diverse (considering the diversity in close relatives), we speculated that mating-type might be a useful proxy for lineage characterization which requires sequence information from only 2-4 genes. As well, categorizing *P. striiformis* f. sp. *tritici* mating-types might give clues as to the evolutionary history of particular *P. striiformis* f. sp. *tritici* lineages, as identifying hybridization events without phased genomic data can be difficult, but the observation of a change in mating-type within an otherwise highly related lineage is a clear indication of a recombination event. Using publicly available sequencing data from over 350 global *P. striiformis* f. sp. *tritici* samples, as well as from 35 Canadian isolates sequenced for this study, we identified nine distinct *P. striiformis* f. sp. *tritici* HD mating-types, and show that mating-type combination is a good proxy for genetic lineage based on whole genome/transcriptome data. Additionally, we connect a recent shift in the northern American *P. striiformis* f. sp. *tritici* population to the appearance of a novel mating-type pair, indicating that the *PstS1*-related lineage of *P. striiformis* f. sp. *tritici* is the product of a recent recombination event between *PstS1* and other existing lineages.

## Results

In order to characterize the history of recombination in the Canadian population of *P. striiformis* f. sp *tritici*, we first identified the set of alleles present at the HD locus across a global dataset. The global dataset consists of 332 previously published RNAseq and gDNA datasets including 17 Canadian samples, and 45 RNAseq datasets derived from samples taken from commercial crop fields in Canada and sequenced in this study. Mating-type genes were identified and characterized using *de novo* assembled transcriptomic data derived from a variety of samples representing diverse lineages of *P. striiformis* f. sp. *tritici* (**Supplementary Data 1**).

A total of nine alleles for each of *Pst-bW-HD1* and *Pst-bE-HD2* were identified, encoding proteins of 593-602 amino acids, and 422-441 amino acids in length, respectively (**Figure 1, Supplementary Data 3**). All alleles encode seemingly functional proteins with the same predicted structure as other basidiomycete HD loci: an N-terminal Variable domain, a central Homeodomain, and a C-terminal Constant domain. The Homeodomain is the only domain with a predicted structure; the other domains are disordered. While the *Pst-bW-HD1* and *Pst-bE-HD2* CDS have an average within-group pairwise nucleotide identity of 80.3% and 78%, respectively, their translated proteins are less well-conserved (77.7% and 75.2%). In all cases the Variable domain has the lowest similarity between any two alleles, usually <60%. The *Pst-bW-HD1* and *Pst-bE-HD2* alleles do not share any meaningful identity outside of the Homeodomain. No indications of recombination between alleles were identified. For example, the *Pst*-*bW2-HD1* allele was always accompanied by a *Pst*-*bE2-HD2* allele, and the same for each other allele pair. *Pst-b1-HD* is unique in that a sub-variant, termed *Pst-b1*-HD* was also identified. The subvariant is identical in the variable region, but contains 12/36 and 17/41 SNP/amino acid polymorphisms in the other domains relative to *Pst*-*bW1-HD1* and *Pst*-*bE1-HD2,* respectively. We are unable to conclude if mating-types *Pst-b1-HD* and *Pst-b1*-HD* are capable of mutual discrimination; however, it seems exceedingly likely given experimental work in *Ustilago maydis* showing that the variable domain is the primary determinant of non-self-recognition [22].

**Figure 1.**
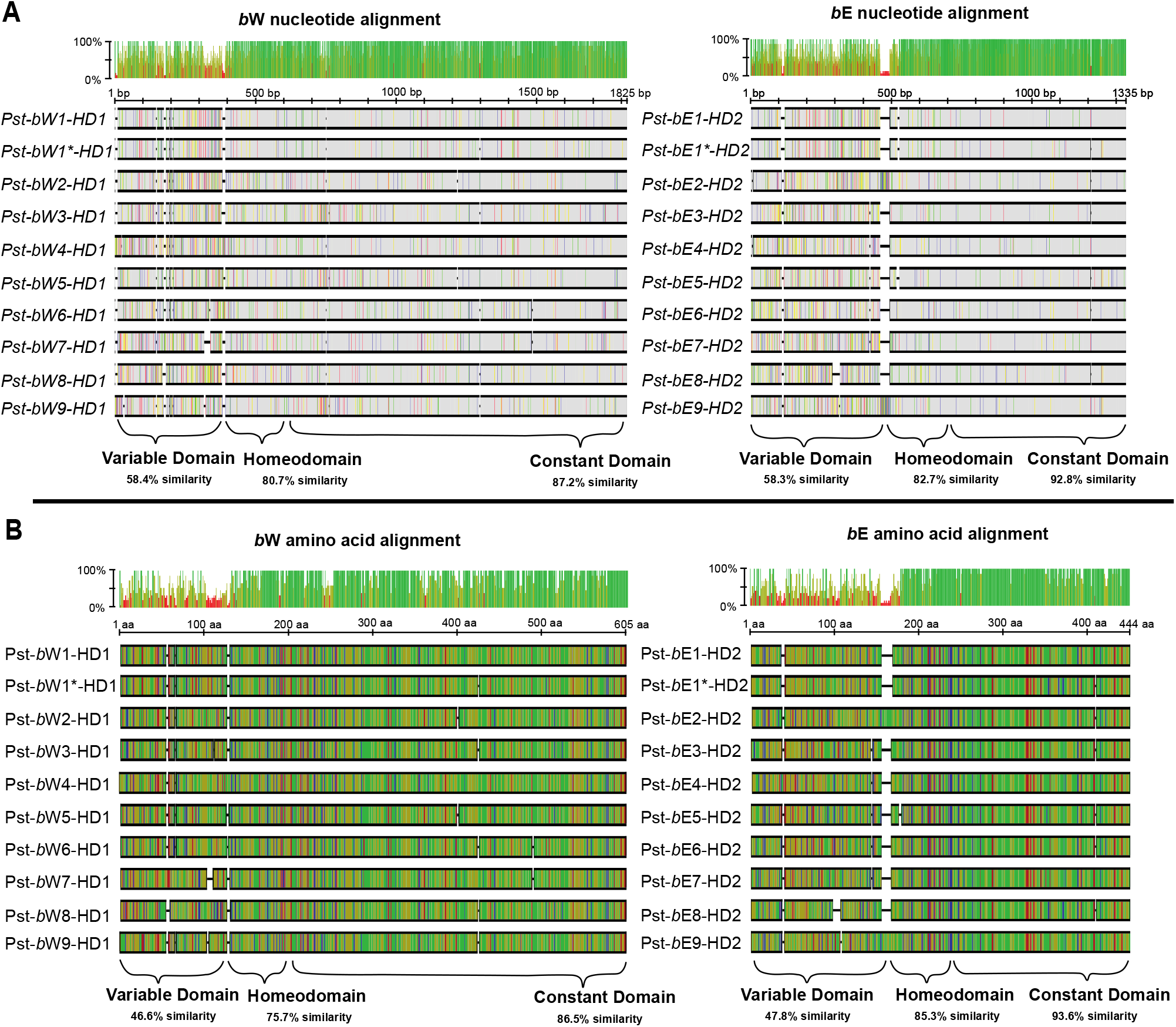
Nucleotide and amino acid diversity across 10 identified *Pst* HD-locus alleles. **(A)** The CDS for each allele of *Pst-bW-HD1* and *Pst-bE-HD2* were aligned using MUSCLE, and visualized in Geneious. Along the top of the alignments is a frequency plot indicating the percentage agreement with the most common nucleotide position. At the bottom of each alignment is the domain structure of the sequences in the alignment, and the average pairwise similarity within that domain across each allele. Each large rectangle represents the nucleotide sequence of a particular allele. Grey stretches indicate agreement with the most common nucleotide at that position. Coloured bars indicate a disagreement, shaded according to the base at that position (Red = A, Green = T, Yellow = G, Blue = C). Alignment gaps due to In/Del polymorphisms between alleles are represented as black horizontal bars, and a scale is provided above the sequences. **(B)** The amino acid alignments of each translated *Pst-bW-HD1* and *Pst-bE-HD2* allele were similarly aligned and cross-compared for agreement at each position. Along the top of the alignments is a frequency plot indicating the percentage agreement with the most common amino acid at that position. Each large rectangle represents the primary sequence encoded by that allele. Amino acids are coloured according to their polarity (Red = D,E; Green = C,N,Q,S,T,Y,U; Orange = A,F,G,I,L,M,P,V,W; Blue = H,K,R.) and gaps in the alignment are indicated by a black horizontal bar. A scale is provided above the sequences.

Having identified the alleles present across this population, we then assessed all available nucleotide datasets for allele presence/absence by searching each dataset for *k*-mers contained within each allele. Nearly all samples (N=345) exhibited *k*-mer signatures for exactly two HD alleles. Datasets containing signatures of more than two alleles (N=18) are believed to be admixtures of more than one isolate, while samples with fewer alleles (N=23) could represent as-yet uncharacterized alleles or simply low sequencing depth of one allele in the dataset (or both).

We also applied this approach to the P/R locus. The STE3 (*Pra*) family of hormone receptor encoding genes and the *mfa* family of hormone precursors have been characterized as determining *­a-locus* specificity. Previous work identified three STE3 family genes in rusts: *STE3.2-1*, *STE3.2-2*, and *STE3.2-3*. *STE3.2-1* and *STE3.2-3* are most closely related to one another and are hypothesized to be biallelic receptor components of this mating locus, which each complement an *mfa*-derived hormone. *STE3.2-2* is thought to be invariant and may be present at a separate genomic locus. We identified all three *STE3.2* genes in these Pst-130 datasets [27], however *STE3.2-2* did not appear in any available transcriptome data, and we conclude that it is not expressed, or not expressed at a high level in either infected leaves or non-germinated urediniospores. *STE3.2-1* and *STE3.2-3* both appeared in the nucleotide data from nearly all isolates (**Supplementary Figure 1**). *STE3.2-2* was detected in genomic data from all isolates in which gDNA sequencing was performed. All *P. striiformis* f. sp. *tritici* samples in this analysis, therefore, share the same biallelic mating-types at the P/R-locus. Isolates where *STE3.2-1* and *STE3.2-3* could not be identified did not contain any other observable STE3 sequence, or any HD sequence, indicating that these samples are likely to represent incompletely sequenced isolates rather than novel mating-type specificities. Interestingly, while *STE3.2-1* and *STE3.2-3* are only 50% similar at the nucleotide level, and both are present in the raw reads of all genomic samples, BLAST search of phased, assembled genomes were only able to identify at most one of these two genes intact, in addition to *STE3.2-2*. Unphased genomes successfully assembled all three *STE3* genes. In no cases were any *STE3* genes identified on the same genomic contig, unlike HD genes which are always found as a pair in head-to-head orientation. Collectively, these results lend credence to the hypothesis that the *STE3.2-1* and *STE3.2-3* genes are part of an allelic series and that their genomic loci may be collapsed by some assemblers.

Having assessed mating-types across the global *P. striiformis* f. sp. *tritici* population, we incorporated this data into a more conventional phylogenomic approach to assessing *P. striiformis* f. sp. *tritici* population structure. In brief, sequence data (RNA and gDNA) was aligned to the reference Pst-130.v2 genome [27], and assessed for intragenic SNPs which were used to construct a maximum-likelihood tree that could be supplemented with HD allele data (**Figure 2**). This analysis was reinforced by using STRUCTURE [28] analysis to identify likely genetic groups from the same intragenic SNP data, and cross-referencing these groups with the clades apparent on the tree. Similar to the work of Radhakrishnan et al. [29], when using a global dataset, STRUCTURE was unable to resolve the more fine-grain distinctions between some closely related sub-clades apparent on the tree. However, taking the first order clades and repeating the analysis successfully, replicated the genetic groups apparent from the phylogeny (**Figure 3**). Groups in our global phylogeny of *P. striiformis* f. sp. *tritici* form into two categories: Clades descended from a single founder isolate, which represent a single, characterized clonal population such as *PstS7/Warrior* or *PstS0*, and population groups which contain a number of related isolates showing signs of admixture and which cannot be said to descend from a single isolate such as isolates sampled in China and Eastern Afrcia/India. Where a group neatly bounds around a previously described clonal lineage, we have annotated the group with that lineage, and where it does not, we have described the origin of the samples within the group. Of note is clade *PstS0*, which represents an extremely old lineage of circulating rusts and so while it can be considered a single clonal population, the individuals within that population exhibit far more intra-clade diversity than, for example, *PstS7/Warrior*. An additional finding of our work is that samples tended to cluster together based on their sequencing manner, i.e., gDNA samples within the *PstS1* clade cluster together relative to RNAseq samples from that same group indicating an unresolved systemic bias in tree construction.

**Figure 2.**
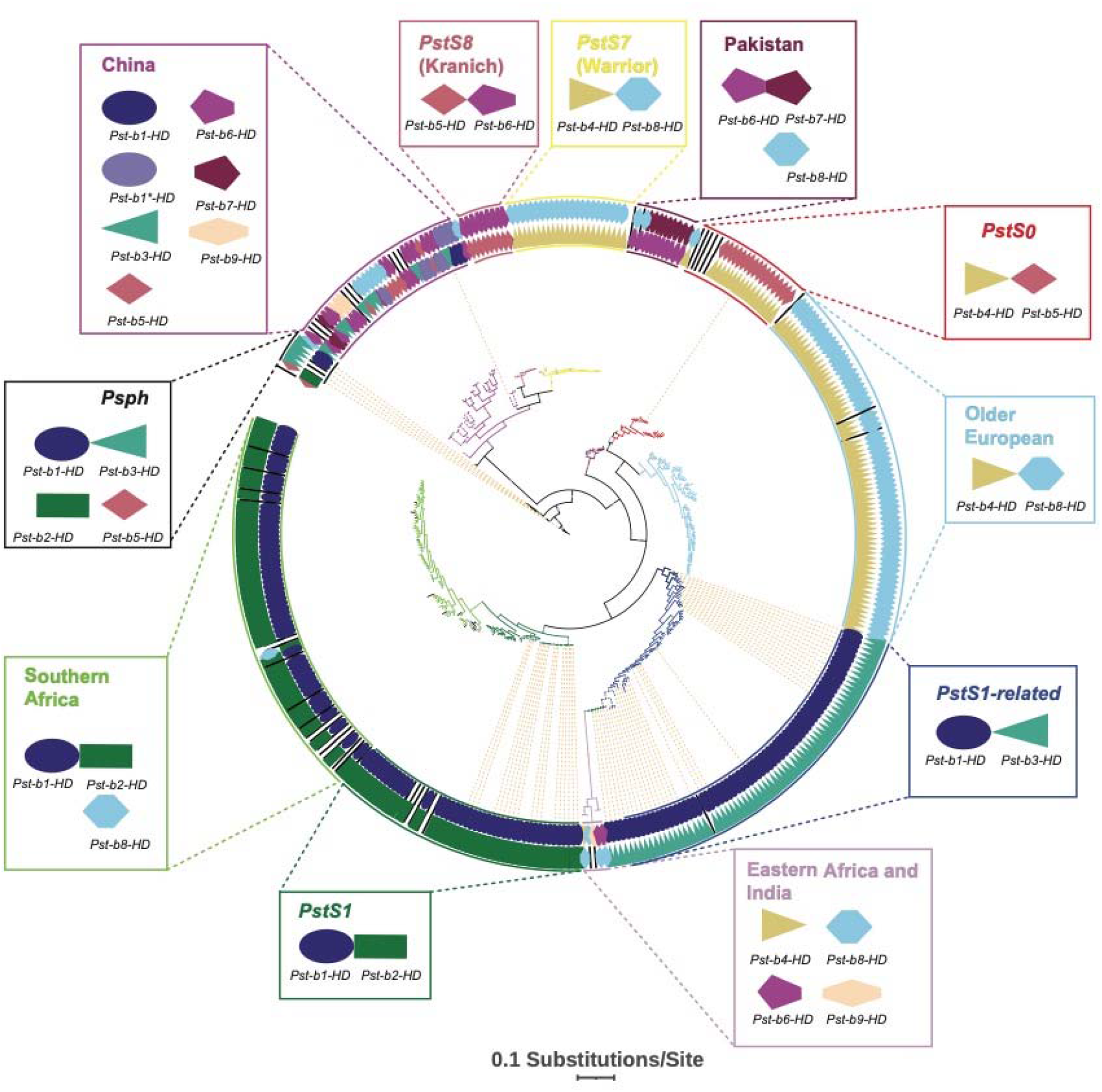
Maximum-likelihood phylogenetic tree detailing the relationship between *Puccinia striiformis* f. sp. *tritici* isolates, annotated with their mating-type alleles. Dendrogram is derived from 131,291 SNPs across all (N=370) datasets. Mating-type alleles present in the isolate are shown as a pair of coloured shapes in the ring surrounding the tree. Unknown mating-types are represented with a black bar. Clades are derived from bootstrap and STRUCTURE analysis and are displayed as coloured branches, matching outer ring colours, and matching text labels. Clade labels are supplemented below with the mating-types identified within the clade. Bootstrap support >80% is shown with a small blue dot at the node. The tree was generated using RAxML and visualised/annotated in IToL and Adobe Illustrator.

**Figure 3.**
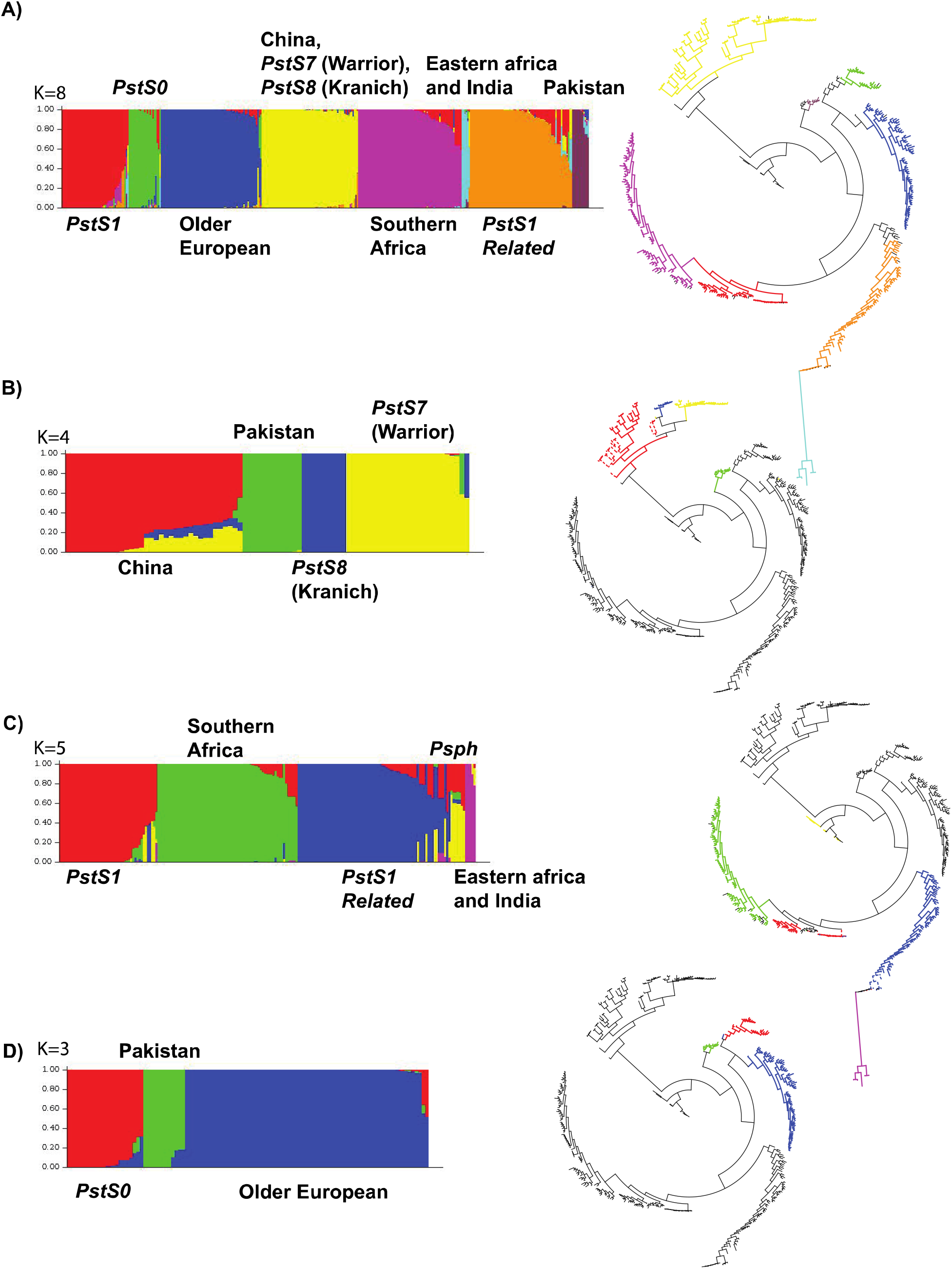
STRUCTURE analyses of the global *Pst* population identified only higher order clades. STRUCTURE analysis of regional *Pst* populations identifies lower order clades. Of the 12 population groups identified by previous work, only seven are identified in a global STRUCTURE analysis, with K>8 not identifying any additional groups. As other groups have reported, sub-analysis of major groups in STRUCTURE was able to correctly identify minor groups. Displayed are the primary, and secondary STRUCTURE analyses, sorted by proposed group (q) and the phylogenetic tree from Figure 2 with the relevant clades highlighted for: (A) All samples (B) Samples from groups 4 and 8 (group 6 hidden for clarity) (C) Samples from groups 1,5,6,7, and *Psph* (D) Samples from groups 2, 3, and 8 (group 6 hidden for clarity).

The majority of the clades worldwide exhibit only a single pair of HD alleles, in addition to short branch lengths and an unbalanced node structure indicating an asexual/clonal population structure where genetic diversity comes from mutations accumulated relative to a single founding isolate (**Figure 2**). European clades both ‘Warrior/Kranich’ (*PstS7/PstS8*) lineages and pre-Warrior, despite forming two robust phylogenetic clades exclusively exhibit the *Pst-b2-HD* + *Pst-b9-HD* mating-type alleles. A single outlying sample exists: Poland 14.0010 which is in the Southern African group and exhibits mating-types *Pst-b2-HD* + *Pst-b8-HD*, rather than *Pst-b1-HD* + *Pst-b2-HD* like its closest relatives (**Supplementary Data 1**). This seems to indicate that recombination (sexual or somatic) in these asexual lineages may infrequently occur but that the resulting lineage will often be outcompeted by its progenitor.

Groups not defined by a single clonal lineage, but rather collected from geographic regions of high diversity and clustering together, specifically those originating in China, eastern Africa and India, and Pakistan exhibit more mating-type diversity with at least seven different HD alleles present in these clades in addition to longer within-group branch lengths and more evenly balanced nodes, indicating that recombination between isolates may be common in those regions. Relative to the large sample size of mostly clonal European and North American populations, it is likely that a substantial portion of *P. striiformis* f. sp. *tritici* diversity in these regions with many different isolates and relatively small sample sizes remains uncaptured.

Having established the distribution of HD locus mating-types across our datasets, and having placed them in the context of the global *P. striiformis* f. sp. *tritici* population, we investigated the North American *P. striiformis* f. sp. *tritici* population specifically. Historically, analyses of North American *P. striiformis* f. sp. *tritici* lineages have grouped Canada and the USA into a single locale, reflecting their large land border and similar climates along each side of the border and due to the fact that stripe rust inoculum in Canada arrives from the USA via the ‘Puccinia pathway’ or wind trajectories along Pacific Northwest [1]. The very first incursion of *P. striiformis* f. sp. *tritici* to north America was in the early 1900s from western Europe and so-called old races belong to the *PstS0* lineage which was restricted to regions west of the Rocky Mountains or southern Alberta. A very small number of isolates from north America were grouped in the *PstS0* clade (**Figure 2**) indicating the lineage is present in north America, which is expected. The other predominant lineage in north America is *PstS1* and several isolates from the USA, Canada, and Kenya were found to be part of this lineage. Prior to 2000, stripe rust was not a disease of concern to growers in Canada (except for southern Alberta), however epidemics in 2000 and 2001 fueled by incursions (from eastern Africa) of races of the *PstS1* lineage to the USA and then to western Canada (via the ‘Puccinia pathway’ along the Great Plains) led to the disease becoming endemic by 2000-2001. Historically, it is believed that the majority of northern American *P. striiformis* f. sp. *tritici* isolates belong to the *PstS1* lineage, with *PstS0* lineage races failing to outcompete this lineage [26,29]. However, contrary to this belief and published literature, the majority of the north American races/isolates after 2015 belong to the distinct *PstS1-related* lineage and not *PstS1. PstS1-related* isolates were only detected in the USA and Canada unlike other lineages which are also present outside north America (**Figure 2**). Other than *PstS0*, *PstS1*, and *PstS1-related* lineages, a single Canadian isolate W056/T210 grouped phylogenetically with the Chinese group of isolates, which is most closely related to the *PstS7/S8* (Warrior/Kranich) lineages. In our previous study [7], W056/T210 was named ‘*PstPr*’ lineage (Pr: probable recombinant) and it was proposed that the lineage is a foreign incursion (due to high telia production ability and close genetic relatedness to *PstS7/S8* isolates). In the present study, we provide conclusive evidence that this race/isolate has its origin in the circulating *P. striiformis* f. sp. *tritici* population within China and is indeed the result of sexual recombination, explaining its high telia production [7].

The HD locus mating-type pair in *PstS1* is *Pst-b1-HD* and *Pst-b2-HD*, but in *PstS1-related* samples the mating-types are *Pst-b1-HD* and *Pst-b3-HD*. It seems likely that the *PstS1-related* clade is the result of fusion between a *PstS1* individual and an individual from another clade with the *Pst-b3-HD* mating-type, possibly belonging to the *PstPr* race along with W056/T210 (*Pst-b1*-HD* and *Pst-b3-HD),* introducing genetic novelty into the north American population and founding a closely related sister-clade to *PstS1*. The *PstPr* lineage was not successful at establishing itself in north America however [7], and no further samples with this unusual configuration have been detected. As of yet, we have insufficient data to evaluate whether this fusion was sexual or somatic by interrogating for synteny between the chromosomes of *PstS1-related* individuals and a *PstS1* individual. The older *PstS1* clade is in fact more closely related to samples taken in the south of Africa than to the *PstS1-related* clade, as evidenced by their positioning in **Figure 2**, their shared mating-types, and STRUCTURE analyses which indicate that while they are distinct populations, they share substantial genetic overlap (**Figure 2, Figure 3**). Indeed, when American and southern African samples are compared directly in STRUCTURE, K=5 identifies the *PstS1, PstS1-related*, southern African, and eastern African and Indian groups as separate clades, while a K>5 continues to describe these same clades but identifies *PstS1* and southern African samples as having a substantially shared genetic background (**Supplementary Figure 2**).

Other than lineages detected in the north American *P. striiformis* f. sp. *tritici* population, our phylogenetic analyses supported a very diverse group of isolates originating mainly in China which the *PstS7* (Warrior) and *PstS8* (Kranich) lineages are derived from, a similar group originating in Pakistan, an unrelated group with samples from India and Eritrea (eastern Africa) including *Pst Race K and Pst Race 21*, and an older western European lineage which appears to be a sister to the *PstS0* lineage in a similar manner to *PstS1* and *PstS1-related*, with each group sharing a single *Pst-HD* allele (*Pst-HD.3*). The final clade, forming an outgroup on the tree is the *Psph* group (*Puccinia striiformis* f. sp. *pseudo-hordei*) which consists of samples collected from foxtail barley grass (*Hordeum jubatum*) as well as the reference genome for *Psh* 93TX-2.

Bayesian STRUCTURE analyses supported the results of our phylogenetic analyses and clustered global *P. striiformis* f. sp. *tritici* population into 8 groups (**Figure 3A, Supplementary Figure 2**). As China lineage, *PstS7*, *PstS8*, and Pakistan lineage appear close to each other, STRUCTURE analyses corroborated their distinction in phylogenetic analyses (**Figure 3B**). Similarly, STRUCTURE supported distinction of *PstS1*, *PstS1-related*, South Africa group, and PstK race at K=5 (**Figure 3C**). *PstS0* and European lineages were also separated by STRUCTURE analyses at K=3 (Figure 3D). It is important to note that *Psph* in the STRUCTURE analyses appears an admixture of multiple lineages from wheat (**Figure 3C**) which is not surprising because the host for *Psph* can harbour rusts from both wheat and barley and helps in rust evolution differently from wheat [7].

It is clear that while mating-types are a critical component of the *P. striiformis* f. sp. *tritici* genome structure and can complement other forms of analyses such as whole genome sequencing, RNAseq and of course phenotyping, they do not fully capture the diversity within a given sample as is made clear by the structures of the *PstS1* and *PstS1-related* groups, as well as the ‘Warrior’ and older European lineages. Given the information provided by assessing the mating-type alleles present across the global population, we were further interested in assessing the diversity within alleles. While some alleles were only present in a few samples, others were present in more than 50 samples spanning over 20 years of data collection, especially those associated with European and northern American agriculture.

## Discussion

Unlike previously published field-pathogenomics studies on *P. striiformis* f. sp. *tritici*, our study placed north American isolates into two distinct lineages with a very clear distinction between *PstS1* and *PstS1-related*, which could be attributed both to a greater number of samples in each clade as well as the fact that we did not rely on a specific subset of genes but took a holistic approach [29,30]. Our approach was to use variation in all coding regions for phylogenetic analyses, the results of which were further supported by independent model-based analyses methods. This also suggest that 242 genes described in Radhakrishnan et al. [29] might not be enough to capture global diversity in the pathogen populations as there are some indications of the division of North American population into two groups in that paper, however, they were not able to conclusively separate the groups into two. A limitation of our approach, however, is the combination of gDNA and RNAseq derived datasets which leads to an observable within-clade segregation between these two origins in the *PstS1* clade, likely due to a combination of systemic error deriving from gDNA and RNAseq reads mapping differently to coding regions as well as differential expression between the two nuclei (i.e., genomic data may capture heterozygous SNPs which are missed in RNAseq data due to a lack of expression).

We identify a shift in the Canadian *P. striiformis* f. sp. *tritici* population after the year 2015 as the predominant lineage changes from *PstS1* to *PstS1-related*. The widespread prevalence of *PstS1-related* isolates over *PstS1* could most likely be attributed to increased fitness in the North American climate, increased urediniospore production or higher aggressiveness on Canadian wheat, but as yet we have no strong evidence for any particular hypothesis. Indeed, in our previous study [7], *PstS1-related* was shown likely to be a recombinant lineage with higher telia production ability than the clonal *PstS1* lineage but the absence of an alternate host in North America should not lead this to favour the *PstS1-related* lineage over *PstS1*. The consequences of *PstS1-related* slowly replacing *PstS1* on Canadian wheat production have not been quantified, but we have not observed a major epidemic due to this incursion, only a single regional epidemic [5] and *PstS1-related* isolates do not appear to exhibit more aggressive or virulent races [data not shown].

This is the first study on wheat stripe rust pathogen *P. striiformis* f. sp. *tritici* to identify and utilize mating-type alleles in pathogenomics and population biology research. Identification of conserved mating-type allele pairs across majority of global lineages further supports the fact that the global population is largely clonal [26,31]. The maximum diversity of mating-type alleles was detected in China which is not surprising as the Himalayan region of China is the centre of origin and diversity of the pathogen [31] and several susceptible barberry (*Berberis* spp.) species as the alternate sexual host of the fungus have been identified from the region where sexual recombination is common [31–35]. From a small number of samples collected from Pakistan and India, there was considerable variation in mating-type alleles and the presence of four alleles in each group suggested some level of sexual recombination which was also reported for isolates from Pakistan in another study [36]. However, the majority of the global lineages originates from a single founder race/isolate and do not show signs of recombination. If recombination is common then replacement of progenitor lineages is uncommon and the emergence of the *PstS1-related* lineage and replacement of *PstS1* as the dominant north American lineage seems to be an unusual event. Continued monitoring of global rust populations taking haplotype and mating-type into account will help to resolve the question of whether hybridization is common and perhaps identify novel lineages as they occur in real-time.

Identification of mating-type alleles can help test biological hypotheses on somatic or sexual recombination in wheat rust pathogen populations. For example, in our previous study [7], the *PstS1-related* lineage was speculated to be a somatic hybrid of *PstS1* and another Canadian lineage, and mating-type allele analysis in the Canadian lineages from this study indicate that the lineage is indeed likely a somatic hybrid between *PstS1* and *PstPr.* To collect further evidence for this claim, our research group is generating phased genome-assemblies of all four lineages from Canada, coupled with chromosome-confirmation, and virulence phenotyping data. Further, we show that mating-type alleles can be used as a proxy for quick lineage identification or prediction, because the majority of the global lineages have unique allele pairs different from other lineages. Adding the HD allele sequences into the MARPLE gene-set used for field-pathogenomics [29] will expand the information to global pathogen population characterization.

## Materials and methods

### Sample collection

In addition to 13 previously published gDNA samples which were included [7], we collected and sequenced the RNA of 43 Canadian rust samples datasets in this study. With the exception of W088 (collected in 1990), all Canadian isolates were collected between years 2005-2021. Additionally, one sample, AR00-05, was collected in 2005 from Arkansas, USA by Dr. Eugene Milus (retired, U. of Arkansas, USA). Samples were collected as leaf tissue infected with a single lesion (single isolated stripe on the leaf) isolates and stored in RNAlater. Such samples are expected to be genetically pure as each successful stripe rust colonization event produces a single stripe along the vascular tissue of the leaf. Isolates described as SP (Single Pustule) have been passed through at least one round of purification through inoculation and spore recovery from a single pustule.

### DNA/RNA extraction and sequencing

Fifty-seven samples were sampled from Canadian fields between 2005 and 2021. Twenty-three were previously used for DNA extraction and WGS [7], and the remaining 34, along with AR00-05 isolate from the USA were used for RNA extraction and sequencing. Eighteen samples were not purified, and RNA was extracted directly from single lesion infected leaf tissue and sequenced with paired-end Illumina technology to a depth of ∼10 Gbp. RNA extraction was performed following protocols described in Radhakrishnan et al. [29]. Seventeen samples were purified to single pustule isolates and RNA was extracted from urediniospores and sequenced with paired-end Illumina technology to a depth of ∼5Gbp.

### Mating-type gene Identification and characterization

In order to characterize the history of recombination in Canadian *P. striiformis* f. sp. *tritici* populations, we first identified the set of alleles present at the HD locus across the global dataset [29,30,37–44]. In the Pst-130.v2 reference genome [27], *Pst-bW-HD1* is represented by FUN_008986+FUN_008987 (partial annotations of a single gene) and by FUN_010468. *Pst-bE-HD2* is represented by FUN_008988 and FUN_010469. Using representatives from each identified clade, we performed *de novo* RNAseq based transcriptome assembly using the Trinity software package with default parameters. As the *Pst-bW-HD1* and *Pst-bE-HD2* genes both possess a conserved Homeodomain and a Constant domain, NCBI Blast+ [45] was used to identify predicted transcripts encoding these genes in the Trinity [46] assembly by querying for the homeodomains identified in the reference genome. Transcripts were then manually curated by aligning the original RNAseq data to the predicted transcripts using BWA [47] and Hisat2 [48], and visualizing the aligned reads in Geneious to curate a biologically plausible pair of alleles for each gene in each isolate (i.e., binning polymorphisms into two alleles based on agreement with paired-end reads. Later, additional samples with unidentified alleles were also passed through this process until no further alleles could be identified.

The *Pst-P/R* complex is represented by the genes FUN_000740 (STE3.2.1), FUN_005623 (STE3.2.2), and FUN_017677 (STE3.2.3). CDS for these genes was extracted and curated with the RNAseq data from isolate W034, to ensure introns had been properly identified. The resulting CDS were carried forward to allele detection in the same manner as the *Pst-HD* locus.

With mating-type alleles identified, the alleles of unknown samples were identified using Sourmash [49] to search for representation of mating-type allele-derived *k*-mers within the raw nucleotide data for that sample. Sourmash sketch was used to create *k*-mer indexes for curated alleles and for nucleotide sequencing data, using the parameters k=21 and 10X downsampling. Sourmash containment with a threshold of 0 was then used to evaluate each sequence dataset for whether or not it carried each allele, the collective outputs are summarized as **Supplementary Data 1**. Samples with over 80% *k*-mer identity for a given allele were generally considered to carry that allele. Unusual samples were manually evaluated for a good match between RNA data and curated mating-types by aligning RNA to the curated mating-type alleles and evaluating the closest match, and in some cases this either prompted curation of another allele, or the allele was left undetermined.

Visualisation of mating-type allele presence by *k*-mers (**Supplementary Figure 1**) was performed using the ggplot2 package in R [50] to generate a heatmap with geom_tile.

### Clade identification and phylogenetic analysis

Thirteen Canadian samples were previously sequenced by the senior author and reported in Brar et al. [7]; the sequence data was utilized in this project. A further 329 global samples were obtained from other previously published studies [27,37–41], in addition to the 44 samples sequenced in this study. Sample origin, mating-type, clade, and other metrics are described in **Supplementary Data 1**.

Illumina reads were aligned to the Pst-130.v2 reference genome using Hisat2 with a minimum score function of L,0,-0.6 and other parameters left default. Alignments were sorted and converted to BAM format using samtools view, and samtools sort [51].

At this stage, quality control was performed on samples by investigating their alignment to the *P. striiformis* f. sp. *tritici* reference genome and to annotated genes within the genome, as well as using a Kraken2 database [52] built from the following preset libraries:

Fungi

Plant

Bacteria

as well as custom libraries constructed from the following genome assemblies:

GCA_001191645.1 *P. triticina*

GCA_000149925.1 *P. graminis*

GCA_001013415.1 *P. arachidis*

GCA_019395275.1 *P. brachypodii*

GCA_002873125.1 *P. coronata*

GCA_002873275.1 *P. coronata*

GCA_008520325.1 *P. graminis*

GCA_007896445.1 *P. hordei*

GCA_001624995.1 *P. horiana*

GCA_004348175.1 *P. novopanici*

GCA_001263375.1 *P. sorghi*

GCA_002920205.1 *P. striiformis*

GCA_019358815.1 *P. triticina*

Of the 17,881 annotated genes in the *P. striiformis* reference genome: average read coverage of >3 reads/bp in fewer than 10,000 genes, low (<20%) *Pucciniales* sample identity in the Kraken2 output, or high (>5%) sample identity from another fungal species in the Kraken2 output were grounds for sample exclusion. 16 samples were excluded this way, described in **Supplementary Data 1**.

For the remaining samples, sorted .bam files were merged into a single mpileup and SNPs called using BCFtools call -m. After calling, SNPs were filtered to intragenic positions using BCFtools filter -R and the set of positions described as exons in the Pst-130.v2 reference GFF3. Following this, informative SNPs were identified using BCFtools filter -i and the parameters: “type==’snp’ && AN >600 && AC/AN>0.01 && AC/AN<0.99 && QUAL>20”, which selected for SNPs in positions where at least 300 isolates had sufficient coverage for a call, overall SNP confidence was >20 (p=0.05), and the minor allele frequency exceeded 0.01. The remaining 131,291 SNPs were converted to phylip format using the scripts at https://github.com/edgardomortiz/vcf2phylip, then used to generate a maximum likelihood tree in RAxML [53] using the following settings:

Mode: -f a

Model: ASC_GTRGAMMA

Bootstraps: 1000

The final tree was visualised with IToL and annotated with clades and mating-type information using iToL and Adobe Illustrator.

### Population STRUCTURE analyses

To delineate the clades described in **Figure 2**, STRUCTURE analyses were performed on the same data. In brief, SNP information was converted from VCF to strct.in format using PLINK [54] and samples were assigned to presumptive clades by cross-referencing for the closest match in Radhakrishnan et al. [29]. STRUCTURE analyses were run using 2000 MCM and burnin reps, and the assumptions of free admixturing and no association between linked markers for values of K between two and 15. The best estimates of K were obtained by comparing the change in ln(Pr|X) between values of K, as well as by investigating the biological plausibility of the resulting groups. In all cases, the point at which increased values of K failed to place any samples into an additional population group agreed with a plateau in ln(Pr|X). After global STRUCTURE comparison was unable to resolve some groups previously identified in other work [28], we performed sub-analyses using related groups identified by the global STRUCTURE analysis. Sub-analyses were able to resolve the missing groups (**Figure 3, Supplementary Figure 2**).

### Data Availability

Sequence data generated in this study and Brar et al. (2018) [7] was uploaded to the NCBI Short Read Archive under bioproject number PRJNA950118.

## Acknowledgements

We are thankful to healthy and fruitful discussions on this work with other wheat rust pathologists in Canada, USA, UK, and Australia.

## Author contributions

**Conceptualization:** Samuel Holden, Guus Bakkeren, Gurcharn S. Brar

**Data curation:** Samuel Holden,

**Formal analyses:** Samuel Holden

**Funding acquisition:** Gurcharn S. Brar, Guus Bakkeren

**Investigation:** Samuel Holden, Gurcharn S. Brar

**Methodology:** Samuel Holden, John Hubensky, Ramandeep Bamrah, Mehrdad Abbasi, Brent D. McCallum, Harpinder S. Randhawa, Muhammad Iqbal, Keith Uloth, Mei-Lan de Graaf, Sang Hu Kim, Rishi Burlakoti

**Project administration:** Gurcharn S. Brar

**Resources:** Dinah Qutob, Hadley R. Kutcher, Gurcharn S. Brar

**Software:** Samuel Holden

**Supervision:** Gurcharn S. Brar

**Validation:** Samuel Holden, Guus Bakkeren, Gurcharn S. Brar

**Visualization:** Samuel Holden

**Writing – original draft:** Samuel Holden, Gurcharn S. Brar

**Writing – review & editing:** Samuel Holden, John Hubensky, Ramandeep Bamrah, Mehrdad Abbasi, Guus Bakkeren, Hadley R. Kutcher, Brent D. McCallum, Harpinder S. Randhawa, Muhammad Iqbal, Keith Uloth, Mei-Lan de Graaf, Sang Hu Kim, Rishi Burlakoti, Gurcharn S. Brar

## Supporting information

**Supplementary Figure S1.**
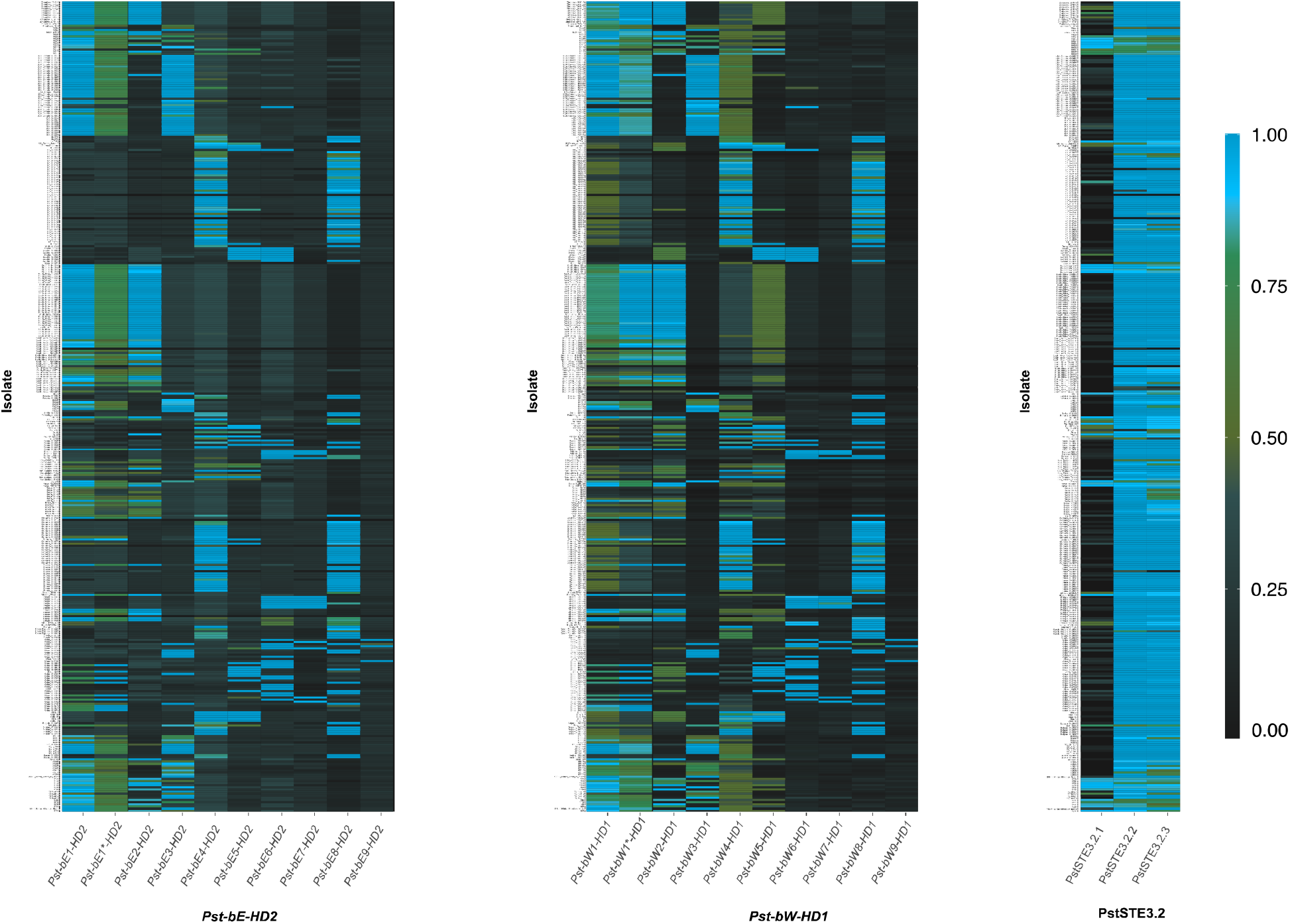
To evaluate the presence or absence of a mating-type allele in a sample, presence of k-mers from that allele was evaluated in each dataset. The percentage of unique k-mers (k=21) from each allele contained in each dataset is rendered as a heatmap, with samples showing >80% presence visualised in light blue, samples with >70% presence in green, and samples with lower k-mer containment in brown trending to black at <50% presence. Sample names are given on the left, and allele names at the bottom. *Pst-b-HD* genes are shown as two groups in the left, and matching alleles are in the same order such that the two genes can be easily compared. *STE3* genes are shown separately on the right, and *STE3.2-2* is only identified in genomic datasets.

**Supplementary Figure S2.**
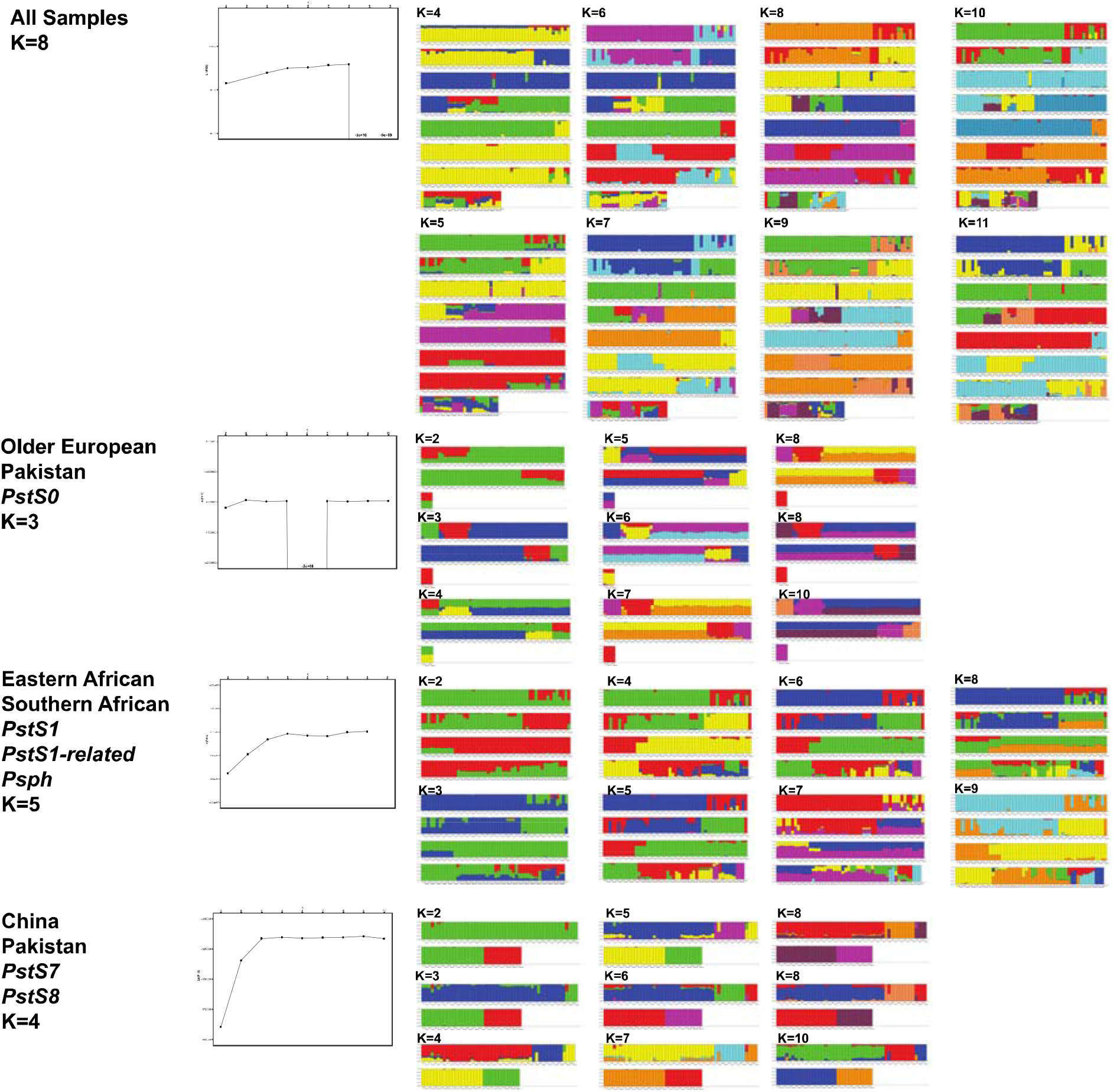
STRUCTURE analysis of the global *Pst* population, as well as previously listed population subgroups for each tested value of K, as well as ln(Pr|X). In this case samples are sorted by ID to enable simple comparison between results for different values of K. Where ln(Pr|X) was radically different from the overall trend, the graph has been supplemented with the value

**Supplementary Data S1. Datasets generated or used in this study.** List of datasets generated or used in this study organised by sample name. Including year of sampling, country of origin, NCBI SRR, assigned clade, and assigned mating-types.

**Supplementary Data S2. Phylogenetic tree displayed in Figure 2, in Newick format**. Tree file includes branch lengths, bootstraps, and isolate names, but not clade information or mating-type annotations. The tree is not rooted in any specific outgroup, and was re-rooted around Psh_93TX-2 in **Figure 2**.

**Supplementary Data S3.** Nucleotide sequence of HD alleles and STE3 genes of *Puccinia striiformis* f. sp. *tritici* identified in this study, in fasta format.

