## Supplementary figures and images for "Uncovering the history of recombination and population structure in western Canadian stripe rust populations through mating-type alleles"

### Supplementary Figure 1

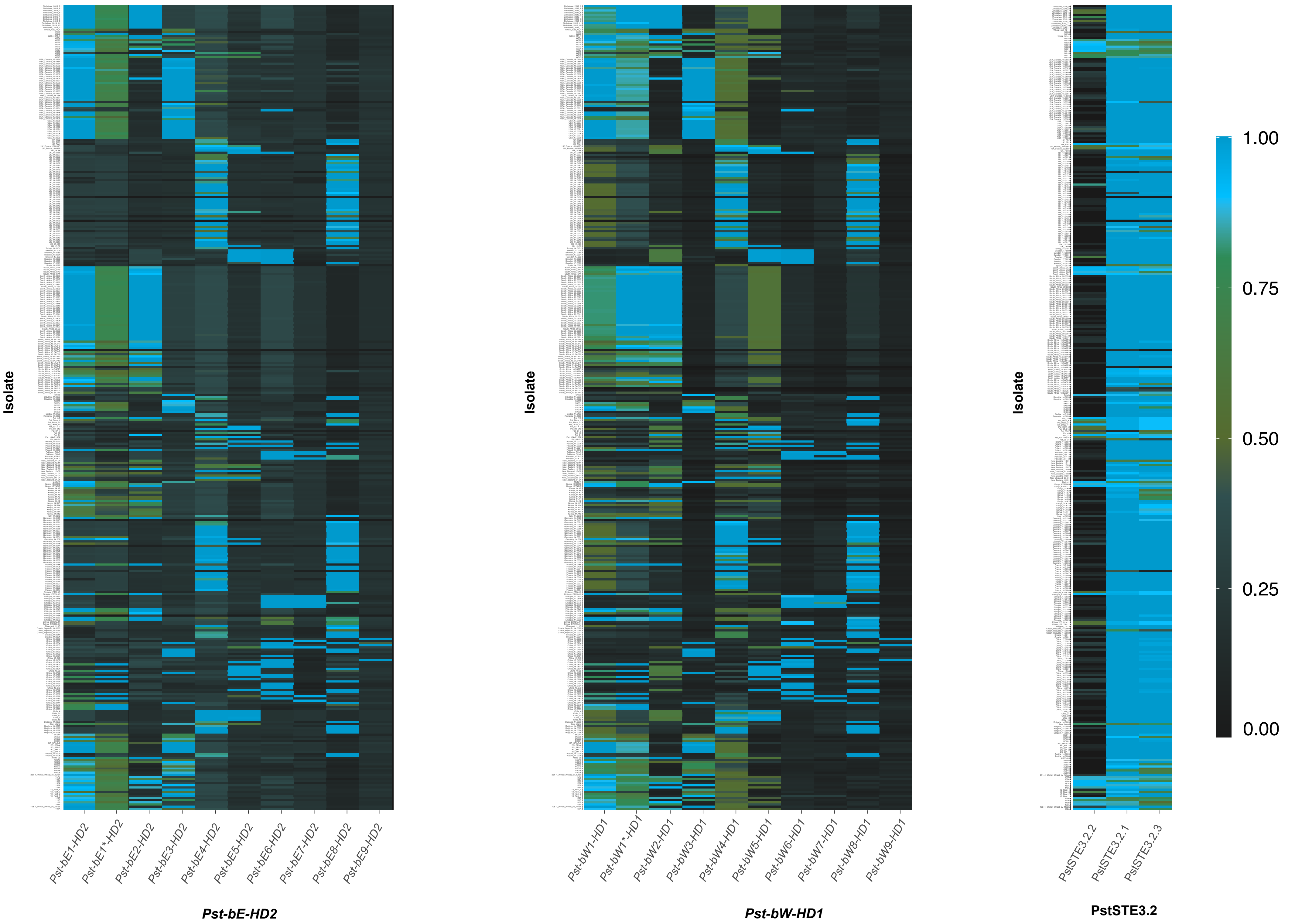

### Supplementary Figure 2

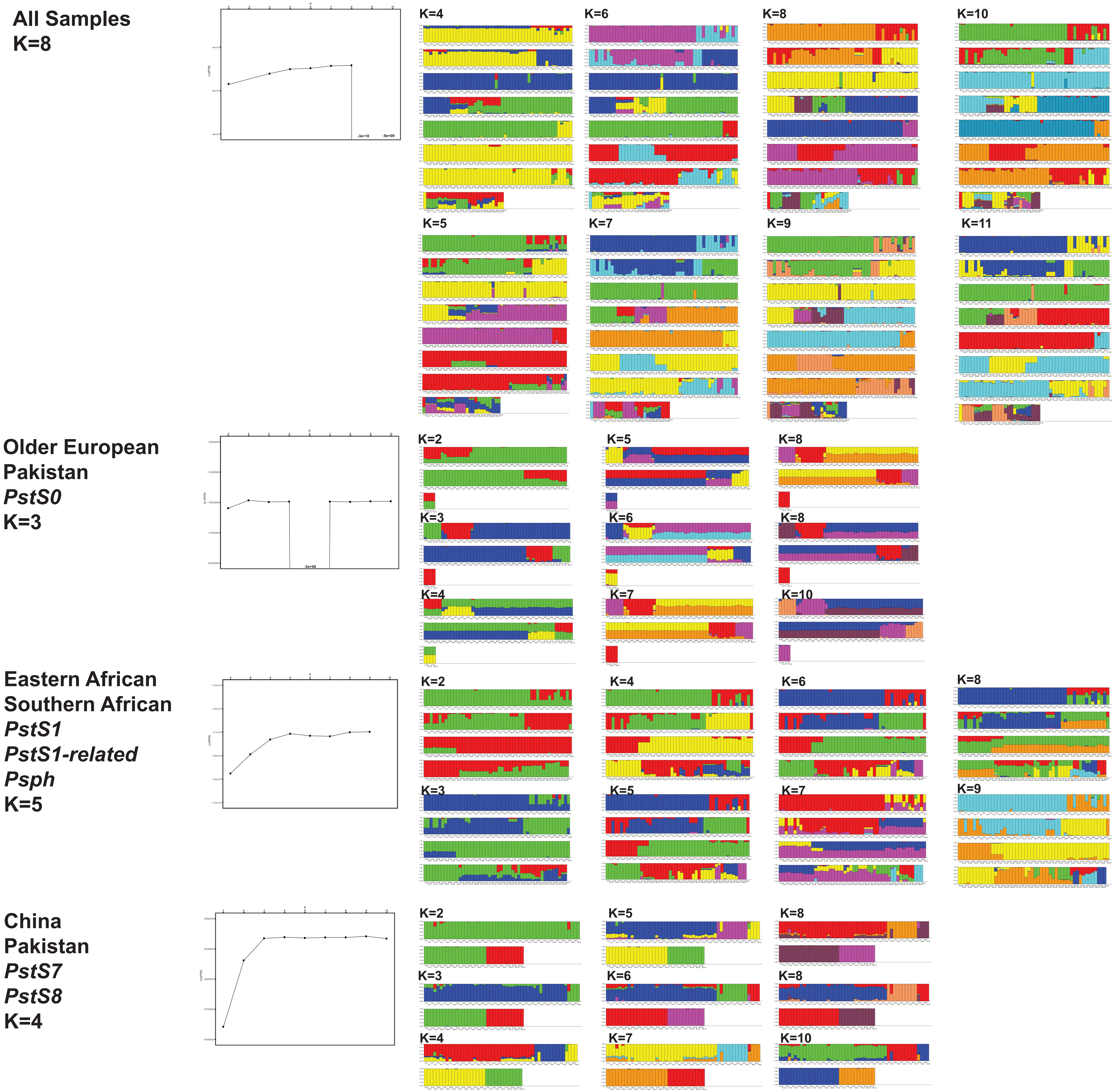
